# A model of compound heterozygous, loss-of-function alleles is broadly consistent with observations from complex-disease GWAS datasets

**DOI:** 10.1101/048819

**Authors:** Jaleal S. Sanjak, Anthony D. Long, Kevin R. Thornton

**Author notes:** (JSS), (KRT).

## Abstract

The genetic component of complex disease risk in humans remains largely unexplained. A corollary is that the allelic spectrum of genetic variants contributing to complex disease risk is unknown. Theoretical models that relate population genetic processes to the maintenance of genetic variation for quantitative traits may suggest profitable avenues for future experimental design. Here we use forward simulation to model a genomic region evolving under a balance between recurrent deleterious mutation and Gaussian stabilizing selection. We consider multiple genetic and demographic models, and several different methods for identifying genomic regions harboring variants associated with complex disease risk. We demonstrate that the model of gene action, relating genotype to phenotype, has a qualitative effect on several relevant aspects of the population genetic architecture of a complex trait. In particular, the genetic model impacts genetic variance component partitioning across the allele frequency spectrum and the power of statistical tests. Models with partial recessivity closely match the minor allele frequency distribution of significant hits from empirical genome-wide association studies without requiring homozygous effect-sizes to be small. We highlight a particular gene-based model of incomplete recessivity that is appealing from first principles. Under that model, deleterious mutations in a genomic region partially fail to complement one another. This model of gene-based recessivity predicts the empirically observed inconsistency between twin and SNP based estimated of dominance heritability. Furthermore, this model predicts considerable levels of unexplained variance associated with intralocus epistasis. Our results suggest a need for improved statistical tools for region based genetic association and heritability estimation.

**Author Summary:** Gene action determines how mutations affect phenotype. When placed in an evolutionary context, the details of the genotype-to-phenotype model can impact the maintenance of genetic variation for complex traits. Likewise, non-equilibrium demographic history may affect patterns of genetic variation. Here, we explore the impact of genetic model and population growth on distribution of genetic variance across the allele frequency spectrum underlying risk for a complex disease. Using forward-in-time population genetic simulations, we show that the genetic model has important impacts on the composition of variation for complex disease risk in a population. We explicitly simulate genome-wide association studies (GWAS) and perform heritability estimation on population samples. A particular model of gene-based partial recessivity, based on allelic non-complementation, aligns well with empirical results. This model is congruent with the dominance variance estimates from both SNPs and twins, and the minor allele frequency distribution of GWAS hits.

## Introduction

Risk for complex diseases in humans, such as diabetes and hypertension, is highly heritable yet the causal DNA sequence variants responsible for that risk remain largely unknown. Genome-wide association studies (GWAS) have found many genetic markers associated with disease risk [1]. However, follow-up studies have shown that these markers explain only a small portion of the total heritability for most traits [2, 3].

There are many hypotheses which attempt to explain the ‘missing heritability’ problem [2–5]. Genetic variance due to epistatic or gene-by-environment interactions is difficult to identify statistically because of, among other reasons, increased multiple hypothesis testing burden [6, 7], and could artificially inflate estimates of broad-sense heritability [8]. Well-tagged intermediate frequency variants may not reach genome-wide significance in an association study if they have smaller effect sizes [9, 10]. One appealing verbal hypothesis for this ‘missing heritability’ is that there are rare causal alleles of large effect that are difficult to detect [4, 11, 12]. These hypotheses are not mutually exclusive, and it is probable that a combination of models will be needed to explain all heritable disease risk [13].

The standard GWAS attempts to identify genetic polymorphisms that differ in frequency between cases and controls. A complementary approach is to estimate the heritability explained by genotyped (and imputed) markers (SNPs) under different population sampling schemes [14, 15]. Stratifying markers by minor allele frequency (MAF) prior to performing SNP-based heritability estimation allows the partitioning of genetic variation across the allele frequency spectrum to be estimated [16], which is an important summary of the genetic architecture of a complex trait [16–23]. This approach has inferred a contribution of rare alleles to genetic variance in both human height and body mass index (BMI) [16], consistent with theoretical work showing that rare alleles will have large effect sizes if fitness effects and trait effects are correlated [18, 20–25]. Yet, simulations of causal loci harboring multiple rare variants with large additive effects predict an excess of low-frequency significant markers relative to empirical findings [4, 26].

SNP-based heritability estimates have concluded that there is little missing heritability for height and BMI, and that the causal loci simply have effect sizes that are too small to reach genome-wide significance under current GWAS sample sizes [14, 16]. Further, extensions to these methods decompose genetic variance into additive and dominance components and find that dominance variance is approximately one fifth of the additive genetic variance on average across seventy-nine complex traits [27]. When taken into account together with results from GWAS, these observations can be interpreted as evidence that the genetic architecture of human traits is best-explained by a model of small additive effects. However, a recent large twin study found a substantial contribution of dominance variance for fourteen out of eighteen traits [28]. The reason for this discrepancy in results remains unclear. One possibility is a statistical artifact; for example, twin studies may be prone to mistakenly infer non-additive affects when none exist. Another possibility, which we return to later, is that this apparently contradictory results are expected under a different model of gene action.

The design, analysis, and interpretation of GWAS are heavily influenced by the ”standard model” of quantitative genetics [29]. This model assigns an effect size to a mutant allele, but formally makes no concrete statement regarding the molecular nature of the allele. Early applications of this model to the problem of human complex traits include Risch’s work on the power to detect causal mutations [30,31] and Pritchard's work showing that rare alleles under purifying selection may contribute to heritable variation in complex traits [17]. When applied to molecular data, such as SNP genotypes in a GWAS, these models treat the SNPs themselves as the loci of interest. For example, influential power studies informing the design of GWAS assign effect sizes directly to SNPs and assume Risch's model of multiplicative epistasis [32]. Similarly, the single-marker logistic regression used as the primary analysis of GWAS data typically assumes an additive or recessive model at the level of individual SNPs [33]. Finally, recent methods designed to estimate the heritability of a trait explained by genotyped markers assigns additive and dominance effects directly to SNPs [14, 16, 27, 34]. Naturally, the results of such analyses are interpreted in light of assumed model of gene action.

A weakness of the multiplicative epistasis model [30, 31] when applied to SNPs is that the concept of a gene, defined as a physical region where loss-of-function mutations have the same phenotype [35], is lost. Specifically, under the standard model, the genetic concept of a failure to complement is a property of SNPs and not ”gene regions” (see [36] for a detailed discussion of this issue). We have recently introduced an alternative model of gene action, one in which risk mutations are unconditionally deleterious and fail to complement at the level of a ”gene region” [36]. This model, influenced by the standard operational definition of a gene [35], gives rise to the sort of allelic heterogeneity typically observed for human Mendelian diseases [37], and to a distribution of GWAS ”hit” minor allele frequencies [4, 26] consistent with empirical results [36]. In this article, we explore this ”gene-based” model under more complex demographic scenarios as well as its properties with respect to the estimation of variance components using SNP-based approaches [34] and twin studies. We also compare this model to the standard models of strictly additive co-dominant effects, and multiplicative epistasis with dominance.

We further explore the power of several association tests to detect a causal gene region under each genetic and demographic model. We find significant heterogeneity in the performance of burden tests [36, 38, 39] across models of the trait and demographic history. We find that population expansion reduces the power to detect causal gene-regions due to an increase in rare variation, in agreement with work by [22, 23]. The behavior of the tests under different models provides us with insight as to the circumstances in which each test is best suited.

In total, our results show that modeling gene action is key to modeling GWAS, and thus plays an important role in both the design and interpretation of such studies. Further, the model of gene-based recessivity best explains the differences between estimates of additive and dominance variance components from SNP-based methods [27] and from twin studies [28] and is consistent with the distribution of frequencies of significant associations in GWAS [4, 26]. Further, the genetic model plays a much more important role than the demographic model, which is expected based on previous work on additive models showing that the genetic load is approximately unaffected by changes in population size over time, [21,22]. Consistent with recent work by [23], we find that rapid population growth in the recent past increases the contribution of rare variants to total genetic variance. However, we show here that different models of gene action are qualitatively different with respect to the partitioning of genetic variance across the allele frequency spectrum. We also show that these conclusions hold under the more complex demographic models that have been proposed for human populations [21, 40].

## Results and Discussion

### The Models

As in [36],we simulate a 100 kilobase region of human genome, contributing to a complex disease phenotype and fitness. The region evolves forward in time subject to neutral and deleterious mutation, recombination, selection, and drift. To perform genetic association and heritability estimation studies *in silico*, we need to impose a trait onto simulated individuals. In doing so, we introduce strong assumptions about the molecular underpinnings of a trait and its evolutionary context.

How does the molecular genetic basis of a trait under natural selection influence population genetic signatures in the genome? This question is very broad, and therefore it was necessary to restrict ourselves to a small subset of molecular and evolutionary scenarios. We analyzed a set of approaches to modeling a single gene region experiencing recurrent unconditionally-deleterious mutation contributing to a quantitative trait subject to Gaussian stabilizing selection. Specifically, we studied three different genetic models and two different demographic models, holding the fitness model as a constant. Parameters are briefly described in Table 1.

**Table 1.** Description of parameters used in the models.

| Parameter | Description |
| --- | --- |
| $N$ | Population size |
| $V_S$ | The total inverse selection intensity |
| $V_E$ | Variance of environmental gaussian deviates |
| $\lambda$ | Mean and standard deviation of trait effects |
| $h$ | Dominance of trait effects |
| $G$ | Genetic contribution to phenotype |
| $E$ | Environmental contribution to phenotype |
| $P$ | Phenotype |
| $w$ | Fitness, based on gaussian function |

We implemented three disease-trait models of the phenotypic form *P* = *G* + *E*. *G* is the genetic component, and 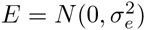 is the environmental noise expressed as a Gaussian random variable with mean 0 and standard deviation 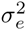. In this context, 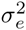 should be thought of as both the contribution from the environment and from the remaining genetic variance at loci in linkage equilibrium with the simulated 100kb region. The genetic models are named the additive co-dominant (AC) model, multiplicative recessive (Mult. recessive; MR) model and the gene-based recessive (GBR) model. The MR model has a parameter, *h*, that controls the degree of recessivity; we call this model the complete MR (cMR) when *h* = 0 and the incomplete MR (iMR) when 0 ≤ *h* ≤ 1. It is important to note that here recessivity is being defined in terms of phenotypic effects; this may be unusual for those more accustomed to dealing directly with recessivity for fitness effects. An idealized relationship between dominance for fitness effects and trait effects of a mutation on an unaffected genetic background is shown in S15 Fig.

The critical conceptual difference between recessive models is whether dominance is a property of a locus (nucleotide/SNP) in a gene or the gene overall. Mathematically, this amounts to whether one first determines diploid genotypes at sites (and then multiplies across sites to get a total genetic effect) or calculates a score for each haplotype (the maternal and paternal alleles). For completely co-dominant models, this distinction is irrelevant, however for a model with arbitrary dominance one needs to be more specific. As an example, imagine a compound heterozygote for two biallelic loci, i.e. genotype Ab/aB. In the case of traditional multiplicative recessivity the compound heterozygote is wild type for both loci and therefore wild-type over all; this implies that these loci are in different genes (or independent functional units of the same gene) because the mutations are complementary. However, in the case of gene-based recessivity [36], neither haplotype is wild-type and so the individual is not wild-type; the failure of mutant alleles to complement defines these loci as being in the same gene [35].

For a diploid with m_i_ causative mutations on the *i*^*th*^ haplotype, we may define the additive model as

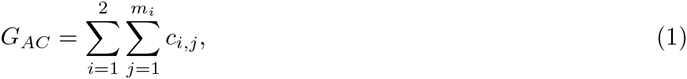

where *c*_*i,j*_ is the effect size of the *j*^*th*^ mutation on the *i*^*th*^ haplotype. Each *c*_*i,j*_ is sampled from an exponential distribution with mean of *λ*, to reflect unconditionally deleterious mutation. In other words, when a new mutation arises it’s effect c is drawn from an exponential distribution, and remains constant throughout it's entire sojourn in the population.

The GBR model is the geometric mean of the sum of effect sizes on each haplotype [36]. We sum the causal mutation effects on each allele (paternal and maternal) to obtain a haplotype score. We then take the square root of the product of the haplotype scores to determine the total genetic value of the diploid.

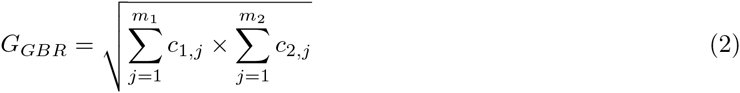

Finally, the MR model depends on the number of positions for which a diploid is heterozygous (*m*_*Aa*_) or homozygous (*m*_*aa*_) for causative mutations,

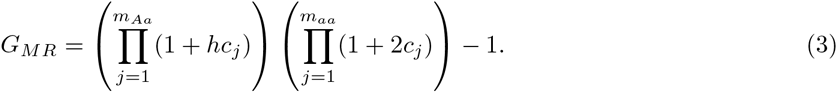

Thus, *h* = 0 is a model of multiplicative epistasis with complete recessivity (cMR), and *h* = 1 closely approximates the additive model when effect sizes are small.

Here, phenotypes are subject to Gaussian stabilizing selection with an optimum at zero and standard deviation of *σ*_*s*_ = 1 such that the fitness, *w*, of a diploid is proportional to a Gaussian function [41].

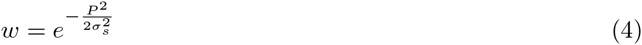

The AC and MR models draw no distinction between a “mutation” and a “gene” (as discussed in [36]). The GBR is also a recessive model, but recessivity is at the level of a haplotype (or allele) and is not an inherent property of individual mutations (see [36] for motivation of this model). Viewed in light of the traditional AC and MR models, the recessivity of a *site* in the GBR model is a function of the local genetic background on which it is found. Based on several qualitative comparisons we find that the GBR model is approximated by iMR models with 0.1 ≤ *h* ≤ 0.25. However, no specific iMR model seems to match well in all aspects. The demographic models are that of a constant sized population (no growth) and rapid population expansion (growth).

The use of the MR model is inspired by Risch’s work [30, 31], linking a classic evolutionary model of multiple loci interacting multiplicatively [42, 43] to the the genetic epidemiological parameter relative risk. Risch and Merikangas [44] used this model to calculate the power detect causal risk variants as a function of their frequency and effect size. Pritchard extended Risch's model to consider a trait explicitly as a product of the evolutionary process [17]. Pritchard's work demonstrated that the equilibrium frequency distribution suggested an important role for rare deleterious mutations when a trait evolves in a constant sized, randomly mating population with recurrent mutation and constant effect sizes. However, multiplicative epistasis is only one model of gene action, and the effect of different genotype-to-phenotype models on the genetic architecture of traits and GWAS outcomes remains unknown. Exploring this issue is the focus of the current work.

### Additive and dominance genetic variance in the population

The amount of narrow sense heritability, *h*^2^ = (*V*_*A*_)/(*V*_*P*_), explained by variants across the frequency spectrum is directly related to the effect sizes of those variants [29]. Thus, this measure is an important predictor of statistical power of GWAS and should inform decisions about study design and analysis [45]. Empirically, SNP-based estimates of heritability have inferred negligible dominance variance underlying most quantitative traits [27]. We have a particular interest in the amount of additive variance, *V*_*A*_, that is due to rare alleles and how much of genetic variance, *V*_*G*_, is attributable to *V*_*A*_ under different recessive models.

We follow the approach of [21], by calculating the cumulative percent of *V*_*G*_ explained by the additive effects of variants less than or equal to frequency *x*, 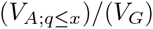. The product of this ratio and broad-sense heritability is an estimate of the narrow-sense heritability, *h*^2^. This calculation is a population-wide equivalent to a SNP-based estimate of heritability in a population sample. In addition we calculate the same distribution for dominance effects 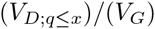 using the orthogonal model of [27]. Methods based on summing effect sizes [29] or the site frequency spectrum [21] would not apply to the GBR model, because the effect of a variant is not independent of other variants (*e.g.*, there is intralocus epistasis). Therefore, we resort to a regression-based approach, where we regress the genotypes of the population onto the total genetic value as defined in our disease trait models (see Material and Methods). In the limit of Hardy-Weinberg and linkage equilibrium, the regression estimates are equivalent to standard quantitative genetic estimates [29] (S14 Fig). For consistency, we applied the regression approach to all models. Overall, these distributions are substantially different across genetic models, demographic scenarios and model parameters (Fig 1).

**Fig 1.**
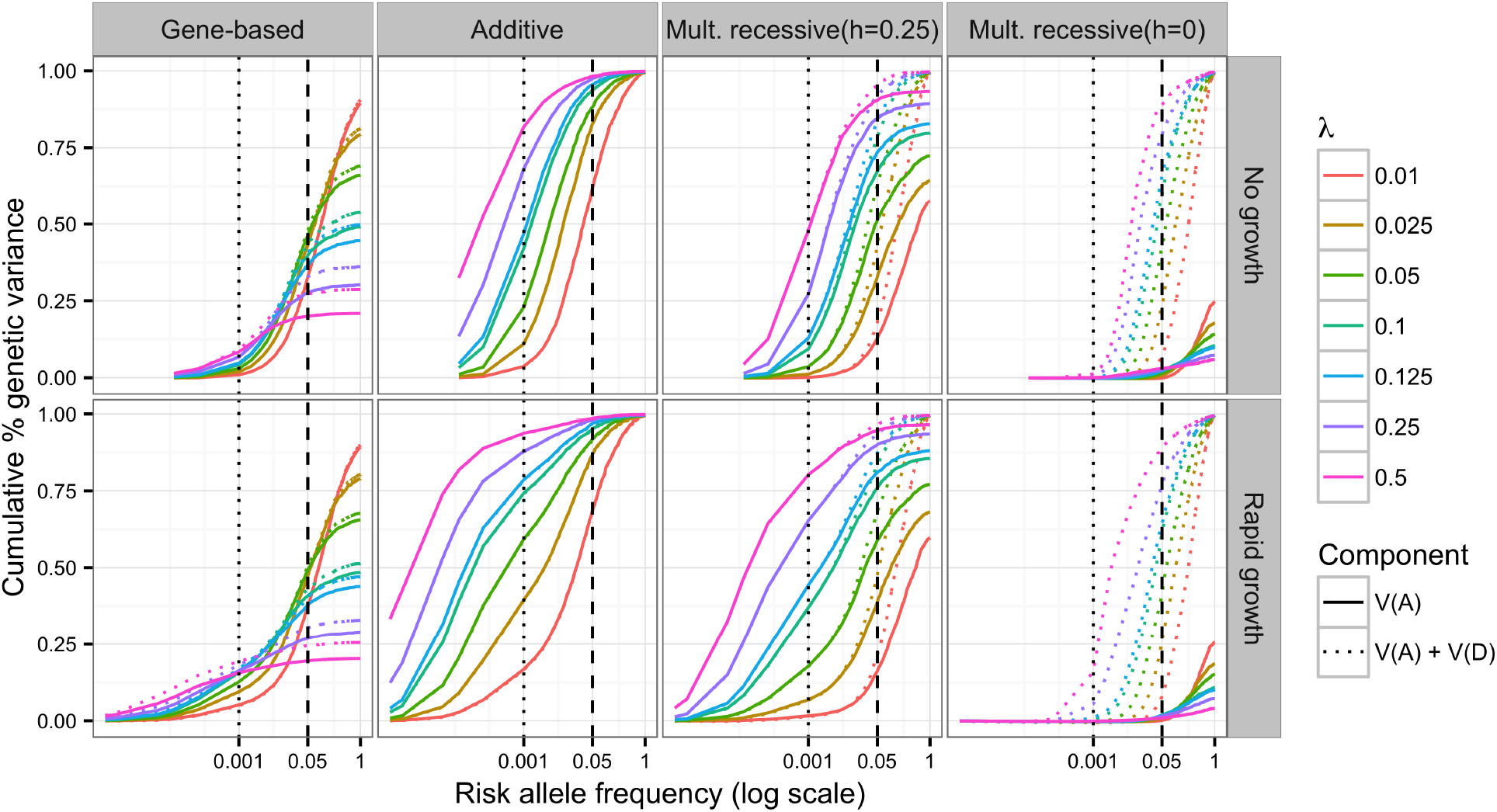
Variance explained over allele frequency. The cumulative additive and dominance genetic variance which can be explained by markers whose frequencies, *q*, are ≤ *x*. Each color represents a different value of *λ*: the mean effects size of a new deleterious mutation. Shown here are the gene-based (GBR), additive co-dominant (AC), incomplete multiplicative recessive (Mult. recessive (*h* = 0.25); iMR) and complete multiplicative recessive (Mult. recessive (*h* = 0);cMR) models. Solid lines show the additive variance alone and dotted lines show the combined additive and dominance variance. All data shown are for models where *H*^2^ ~ 0.08, although it was confirmed (data not shown) that these particular results are independent of total *H*^2^. The additive and dominance genetic variance is estimated by the adjusted *r*^2^ of the regression of all markers (and their corresponding dominance encoding) with *q* ≤ *x* onto total genotypic value (see methods for details); data are displayed as the mean of 250 simulation replicates. The vertical dotted and dashed lines correspond to the *q* = 0.001 and *q* = 0.05, respectively. The curves under no growth appear to be truncated with respect to rapid growth because the range of the x-axis differs between growth and no growth (minimum *q* = 1/2*N*).

Under the AC model, all of *V*_*G*_ is explained by additive effects if all variants are included in the calculation; in Fig 1 the solid variance curves reach unity in the AC panel. Low frequency and rare variants (*q* < 0.01) explain a large portion of narrow sense heritability (26% – 95%) even in models without rapid population expansion. Further, the variance explained at any given frequency threshold increases asymptotically to unity as a function of increasing *λ* (S4 Fig). While the total heritability of a trait in the population is generally insensitive to population size changes (S1 Fig, see also [21, 22, 46]), rapid population growth increases the fraction of additive genetic variation due to rare alleles (Fig 1).

Here, increasing *λ* corresponds to stronger selection against causative mutations, due to their increased average effect size. Recent work by Zuk et al. [24], takes a similar approach and relates the allele frequency distribution directly to design of studies for detecting the role of rare variants. However, our findings contrast with those of Zuk [24] and agree with those of Lohmueller [22], in that we predict that population expansion will substantially increase the heritability, or portion of genetic variance, that is due to rare variants. Our results under the AC model agree with those of Simons et al. [21], in that we find that increasing strength of selection, increasing *λ* in our work, increases the contribution to heritability of rare variants. However, under the GBR model and the cMR model the distribution of genetic variance over risk allele frequency as function *λ* is non-monotonic (Fig 1 and S4 Fig).

For all recessive models, we find that total *V*_*A*_ is less than *V*_*G*_ (Fig 1). For the MR models, all additional genetic variation is explained by the dominance variance component; in Fig 1 the dotted variance curves reach unity in the MR panels. As expected, genetic variation under the MR model with partial recessivity (*h* = 0.25) is primarily additive [29, 47], whereas *V*_*G*_ under the cMR model (*h* = 0) is primarily due to dominance. The GBR model shows little dominance variance and is the only model considered here for which the total *V*_*G*_ explained by *V*_*A*_ + *V*_*D*_ is less than the true *V*_*G*_ for all *λ*. This can be clearly seen in Fig 1 where the dotted curves do not reach unity in the GBR panel. These observations concerning the GBR model are consistent with the finding of [27] that dominance effects of SNPs do not contribute significantly to the heritability for complex traits.

Under the GBR model, large trait values are usually due to compound heterozygote genotypes (*e.g.*, *Ab/aB*, where A and B represent different sites in the same gene) [36]. Therefore, the recessivity is at the level *of the gene region* while the typical approach to estimating *V*_*A*_ and *V*_*D*_ assigns effect sizes and dominance to individual mutations. Thus, compound heterozygosity, which is commonly observed for Mendelian diseases (see [36] and references therein) would be interpreted as variation due to *interactions* (epistasis) between risk variants. Importantly, the GBR model assumes that such interactions should be local, occurring amongst causal mutations in the same locus. While the GBR model is reflective of the original definition of a gene in which recessive mutations fail to complement, we emphasize that this does not imply that mutations are necessarily exomic. The GBR model is of a general genomic region in which mutations act locally in *cis* to disrupt the function of that region with respect to a phenotype.

The increase in the number of rare alleles due to population growth is a well established theoretical and empirical result [48–61]. The exact relationship between rare alleles [4, 17, 26, 62, 63], and the demographic and/or selective scenarios from which they arose [21, 22, 64], and the genetic architecture of common complex diseases in humans is an active area of research. An important parameter dictating the relationships between demography, natural selection, and complex disease risk is the degree of correlation between a variant’s effect on disease trait and its effect on fitness [18, 20–22]. In our simulations, we do not impose an explicit degree of correlation between the phenotypic and fitness effects of a variant. Rather, this correlation is context dependent, varying according to the current genetic burden of the population, the genetic background in which the variant is present and random environmental noise. However, if we re-parameterized our model in terms of [18], then we would have *τ* ≤ 0.5 (Gaussian function is greater than or equal to its quadratic approximation), which is consistent with recent attempts at estimating that parameter [20, 65]. Our approach is reflective of weak selection acting directly on the complex disease phenotype, but the degree to which selection acts on genotype is an outcome of the model. While the recent demographic history has little effect on key mean values such as broad-sense heritability of a trait or population genetic burden (S1 Fig and S3 Fig), the structure of the individual components in the population which add up to those mean values varies considerably. The specific predictions with respect to the composition of the populations varies drastically across different modeling approaches. It is therefore necessary to carefully consider the structure of a genetic model in a simulation study.

The conclusions reached here also hold when we consider more complex demographic scenarios relevant to human populations. Under the demographic model for European populations from [40], the additive and GBR models show the same behavior as in Fig 1 (S17 Fig). At all key time points where population size changes, *V*_*A*_ = *V*_*G*_ for the additive model, and the variance explained by rare mutations depends primarily on *λ* (S17 Fig). For the GBR model, *V*_*A*_ < *V*_*G*_ (as in Fig 1), and plateaus at the same ratio *V*_*A*_/*V*_*G*_ for all time points except immediately after the bottleneck, which results in a short-lived increase in *V*_*A*_/*V*_*G*_ that is undetectable by the time growth begins (S17 Fig). All recessive models (GBR, iMR and cMR) may show a transient increase in total *V*_*G*_ after the bottleneck, depending on the value of *λ* (S18 Fig). However, the GBR and iMR models with *h* > 0.25 showed a return to constant population size levels by the final time point. The changes in *V*_*A*_ and *V*_*G*_ under recessive models is likely due to the transfer of non-additive variation into *V*_*A*_ during a bottleneck, which has been studied thoroughly in the theoretical literature [66, 67]. As in Fig 1, the genetic model, and not the demographic details, drive the relationship between mutation frequency and additive genetic variance.

### Estimating additive and dominance variance from population samples

The previous section shows that the relationship between genetic variance and allele frequency in the entire population strongly depends on the genetic model. Recent estimates of variance components from large population samples of unrelated individuals have inferred that dominance variance (*V*_*D*_) is negligible for most traits [27]. However, a recent study of more than 10^4^ Swedish twins and 18 traits obtained a contradictory result, inferring significant non-additive variance for most traits, which was interpreted as *V*_*D*_ [68]. In this section, we show that this apparent inconsistency is expected under certain models of gene action.

We applied GREMLd, MAF-stratified GREMLd (MS-GREMLd), and MAF-stratified Haseman-Elston regression (see Methods for details). We found MS-GREMLd to be numerically unstable on our simulated data, and thus we present results for non-MS-stratified GREMLd. The numerical stability issues likely resulted from some combination of small number of SNPs per region 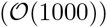, low total *V*_*G*_ in a region, or high variance in effect sizes across causal mutations [69]. Further, for large *λ*, where *V*_*G*_ is primarily due to rare alleles (Fig 1), heritability in a sample may not reflect heritability in the entire population (S13 Fig).

Fig 2 shows the GREMLd additive and dominance heritability estimates, as compared to the respective population value, over *λ*. Under the cMR model (*h* = 0), the dominance component is much larger than the additive component as predicted from Fig 1. When GREMLd is performed on data after removing variants with *MAF* ≤ 0.01, as done in [27], the total heritability estimate (AD) is quite accurate. As anticipated, GREMLd using unfiltered data yields results with a slight upward bias [70]. However, for the iMR (*h* = 0.25) model the filtered GREMLd estimates are only accurate for *λ* < 0.1 reflecting the preponderance of rare causal variants for larger values of *λ*. Unfiltered GREMLd estimates under the iMR (*h* = 0.25) model show a slight upward bias for small values of *λ*, but are otherwise accurate. This shows that GREMLd is performing as expected under the site-based model for which it is designed. The MS-HE regression results are generally consistent with the GREMLd results.

**Fig 2.**
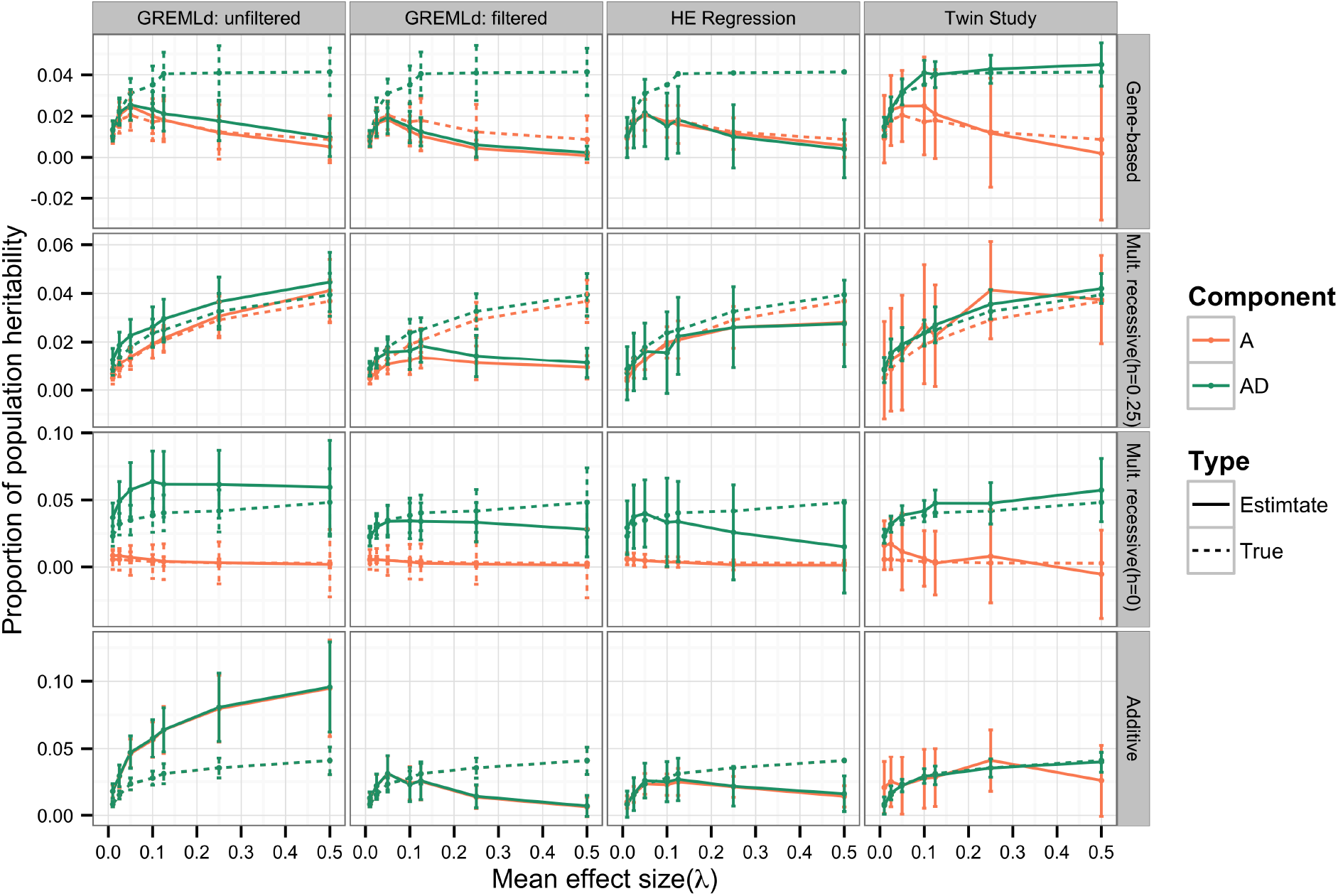
Heritability estimates compared to population heritability. Heritability estimates and population heritability as a function of *λ*: the mean effect size of a new deleterious mutation. Additive (A; orange) component of true heritability is calculated by multiplying the end point(*q* = 1) of the variance curves in Fig 1 by the broad-sense heritability values summarized in S1 Fig. HE-regression and GREMLd estimates were obtained from random population samples (n = 6000). GREMLd analysis was performed in GCTA using genotype data that was either unfiltered or filtered to remove variants with MAF<0.01. Twin study estimates are directly calculated using MZ and DZ twin correlations from 64 sets of twin studies. Each study consisted of pooling 2000 MZ twin pairs and 2000 DZ twin pairs from each of 8 model replicates for a total of 64,000 individual phenotypes. Data are plotted as the median across replicate sets ± half the interquartile range. Shown are the additive co-dominant (AC), gene-based (GBR) incomplete multiplicative recessive (Mult. recessive (*h* = 0.25); iMR) and complete multiplicative recessive (Mult. recessive (*h* = 0); cMR) models.

The GREMLd and MS-HE estimates are accurate under the GBR model when *λ* is small, because most heritability is additive in that case(Fig 1). However, under the GBR model, both filtered and unfiltered GREMLd heritability estimates show downward bias when *λ* is large (Fig 2). The MS-HE regression results reveal a similar pattern, which indicates that the downward bias for large values of *λ* is not strictly due to removal of rare variants in the filtered GREMLd analysis. Instead, the bias shown for large values of *λ* is likely due to the presence of substantial non-additive heritability, which is not captured by the dominance effects of SNPs.

In contrast to the variance component methods, our simulated large twin studies provide approximately unbiased estimates of total heritability for large values of *λ*, but were biased upward for small effect sizes under the AC and GBR models (Fig 2). The variance in twin-study estimates was large enough that study-to-study disagreement would be expected. Formally, twin studies estimate an additive and a non-additive component of variance and interpreting the non-additive component as epistatic or dominance variance is a matter of perspective. However, the GBR model is inspired by the definition of a gene as a physical region in which recessive mutations leading to the same phenotypic outcome fail to complement [35], consistent with the allelic heterogeneity observed for human Mendelian disorders (see [36] for further discussion). Thus, the model of recessivity at the level of the gene region is picked up as non-additive variance in twin studies, but missed by variance component methods (GREML and HE regression) because the dominance in the GBR model is due to *Ab/aB* (compound heterozygotes) genotypes rather than *a/a* genotypes (heterozygotes for a specific loss of function variant) assumed by variance component methods. Thus the contradictory results of applying variance component methods [27] and analysis of large twin studies [68] in order to estimate *V*_*A*_ and *V*_*D*_ may be interpreted as evidence for a model of gene action such as the GBR, which may be viewed as either recessivity at the haplotype/gene level or intralocus epistasis at the level of causative mutations in a single gene region. Both interpretations are valid. The alternative explanation is that we must assert that one of the study designs is generating artifacts.

### The genetic model affects the outcomes of GWAS

Both demography and the model of gene action affect the degree to which rare variants contribute to the genetic architecture of a trait (Fig 1). However, the different mappings of genotype to phenotype from model to model make it difficult to predict *a priori* the outcomes of GWAS under each model. Therefore, we sought to explicitly examine the performance of statistical methods for GWAS under each genetic and demographic model. We assessed the power of a single marker logistic regression to detect the gene region by calculating the proportion of model replicates in which at least one variant reached genome wide significance at *α* ≤ 10^−8^ (Fig 3A). The basic logistic regression is equivalent to testing for association under the AC model. We simulated both a perfect “genotyping chip” (all markers with MAF ≥ 0.05) and complete re-sequencing including all markers (Fig 3B).

**Fig 3.**
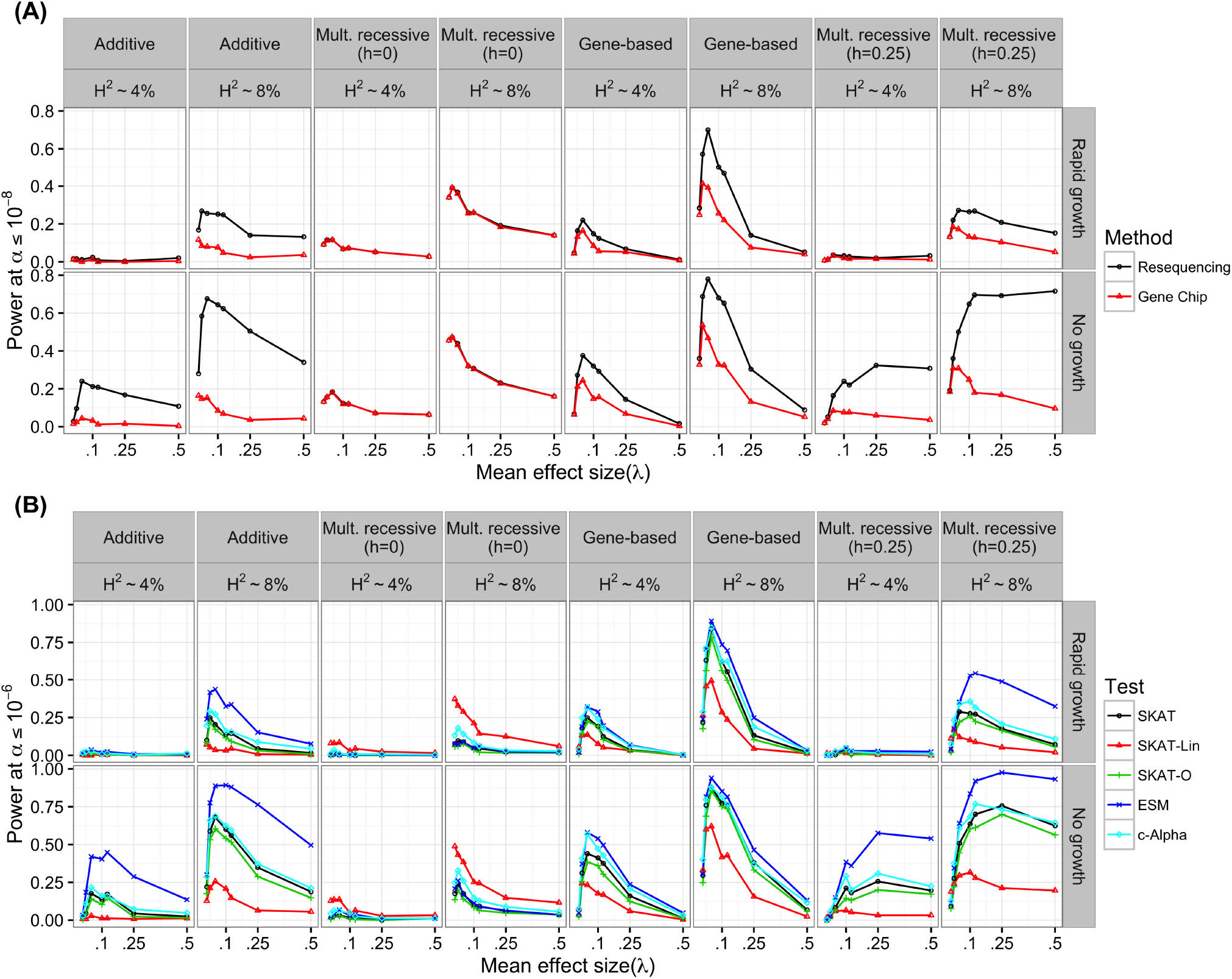
Power of association tests. (A) The power of a single marker logistic regression, at significance threshold of *α* ≤ 10^−8^, as a function of *λ*: the mean effect size of a new deleterious mutation. For single marker tests we define power as the number of simulation replicates in which any single marker reaches genome wide significance. Two study designs were emulated. For the gene chip design only markers with *MAF* > 0.05 were considered and all markers were considered for the resequencing design. Genetic models shown here are the additive co-dominant (AC), gene-based (GBR), complete multiplicative site-based recessive (Mult. recessive (*h* = 0); cMR) and incomplete multiplicative site-based recessive models (Mult. recessive (*h* = 0.25); iMR) (B) The power of region-based rare variant association tests to detect association with the simulated causal gene region at significance threshold of *α* ≤ 10^−6^. For region-based tests, we define power as the percent of simulation replicates in which the p-value of the test was less than *α*. The p-values for the ESM, c-Alpha were evaluated using 2 × 10^6^ permutations. SKAT p-values were determined by the SKAT R package and represent numerical approximations to the presumed analytical p-value.

Across all genetic models, the single marker logistic regression has less power under population expansion (Fig 3A). The loss of power is attributable to a combination of rapid growth resulting in an excess of rare variants overall [48–61], and the increasing efficacy of selection against causal variants in growing populations [21]. While complete resequencing is more powerful than a gene-chip design, the relative power gained is modest under growth (Fig 3A). Region-based rare variant association tests behave similarly with respect to population growth (Fig 3B).

There are important differences in the behavior of the examined statistical methods across genetic models. We focus first on the single marker tests (Fig 3A). For gene-chip strategies, power increases for “site-based” models as recessivity of risk variants increases (compare power for AC, iMR, and cMR models in Fig 3B). This increase in power is due to the well-known fact that recessive risk mutations are shielded from selection when rare (due to being mostly present as heterozyogtes), thus reaching higher frequencies on average (S5 Fig), and that the single-marker test is most powerful when risk variants are common [32]. Further, for the complete multiplicative-recessive model (cMR), the majority of *V*_*G*_ is due to common variants (Fig 1), explaining why resequencing does not increase power for this model (Fig 3A).

For single-marker tests, the GBR model predicts large gains in power under re-sequencing for intermediate *λ* (the mean trait-effect size of newly arising causal mutations), similar to the AC or iMR model. But, when *λ* is larger power may actually be less under the GBR model than under AC or iMR. For all models, causal mutations are more rare with increasing lambda (S7 Fig). However, as a function of frequency, all *V*_*G*_ may be attributed to *V*_*A*_ or *V*_*D*_ in the site-specific models whereas there is increasing intralocus epistasis in the GBR model as a function of *λ* (Fig 1). It is well-known that the single marker test has lower power when causal mutations have low frequencies, are poorly tagged by more common SNPs, or have small main effects [32, 71].

Region-based rare variant association tests show many of the same patterns across genetic model and effect size distribution as single marker tests, but there are some interesting differences. The ESM test [36, 72] is the most powerful method tested for the AC, iMR, and GBR models (Fig 3b), with the c-Alpha test as a close second in some cases. For those models, the power of naive SKAT, linear kernel SKAT and SKAT-O, is always lower than the ESM and c-Alpha tests. This is peculiar since the c-Alpha test statistic is the same as the linear kernel SKAT test. The major difference between SKAT and ESM/c-Alpha is in the evaluation of statistical significance. SKAT uses an analytical approach to determine p-values while the ESM/c-Alpha tests use an explicit permutation approach. This implies that using permutation based p-values results in greater power. Yet, under the cMR model the linear kernel SKAT is the most powerful, followed by c-alpha. The cMR model does not predict a significant burden of rare alleles and so the default beta weights of SKAT are not appropriate, and the linear kernel is superior. The ESM test does poorly on this model because there are not many marginally significant low-frequency markers. It is logical to think that these tests would all perform better if all variants were included. The massive heterogeneity in the performance of region-based rare variant tests across models strongly suggests that multiple methods should be used when prior knowledge of underlying parameters is not available. In agreement with [22, 73], we predict that population growth reduces the power to associate variants in a causal gene region with disease status (Fig 3) when the disease also impacts evolutionary fitness. We have recently released software to apply the ESM test to case control data [72] in order to facilitate applying this test to real data.

### The distribution of minor allele frequencies of GWAS hits

It was noted by [4, 26], that an excess of rare significant hits, relative to empirical data, is predicted by AC models where large effect mutations contribute directly to fitness and the disease trait. We confirm that AC models are inconsistent with the empirical data (Fig 4), except when *λ* ≤ 0.01. The empirical data in Fig 4 represent a pooled data set with the same diseases and quality filters as in [26], but updated to include more recent data. The data are described in S1 Table, and can be visualized alone more clearly in S16 Fig. Close to half of the data comes from GWAS studies uploaded to the NHGRI database after 2011, yet the same qualitative pattern is observed. This contradicts the hypothesis that the initial observation of an excess of common significant hits relative to the prediction under an AC model was simply due to small sample sizes and low marker density in early GWAS previously analyzed in [4, 26]. Yet the initial observation is in fact robust and the meta-pattern provides an appropriate point of comparison when considering the compatibility of explicit population-genetic models with existing GWAS data.

**Fig 4.**
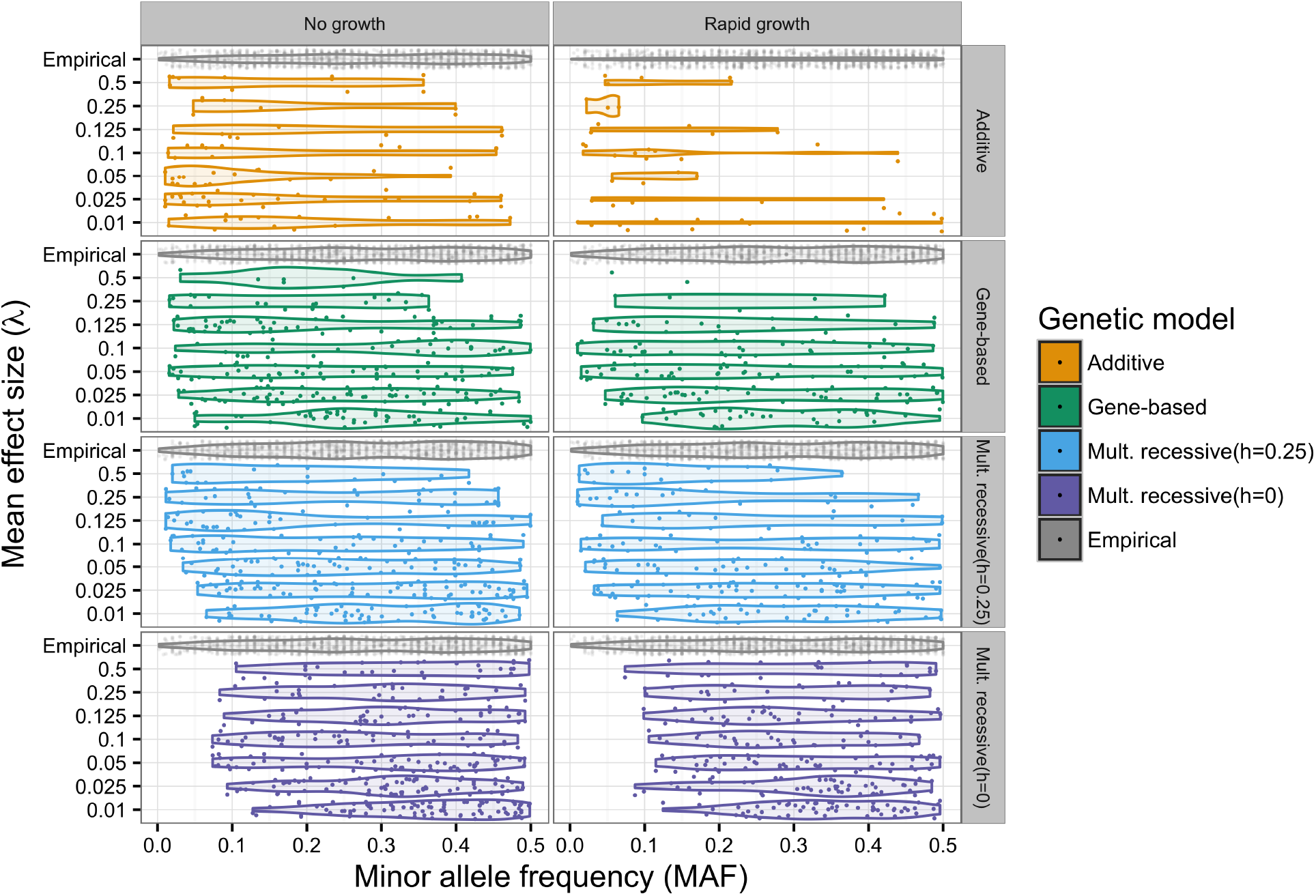
Distribution of significant GWAS hits. Horizontal violin plots depict the distribution of minor allele frequencies (MAF) of the most strongly associated single marker in a GWAS. Individual hits are plotted as translucent points and jittered to provide a sense of the total number and density of hits. Each panel contains simulated data pooled across model replicates for each value of *λ* with empirical data for comparison. Empirical data are described in Materials and Methods. In cases where more than one marker was tied for the lowest p-value, one was chosen at random. Shown here are the additive co-dominant (AC), gene-based (GBR), incomplete multiplicative recessive (Mult. recessive (*h* = 0.25); iMR) and complete multiplicative recessive (Mult. recessive (*h* = 0);cMR) models. All data shown are for models where *H*^2^ ~ 0.08, because single marker test power was too low under *H*^2^ ~ 0.04 to make informative density plots. Simulated data were subjected to ascertainment sampling such that the MAF distribution of all markers on the simulated genotyping chip was uniform. Specific information regarding the empirical data can be obtained in S1 Table.

The GBR model predicts few rare significant hits and an approximately uniform distribution across the remainder of MAF domain (Fig 4), even for intermediate and large values of *λ*. For smaller values of *λ*, the GBR predicts an excess of common significant hits. The more uniform distribution of significant single markers seen under the GBR is consistent with the flatter distribution of genetic variance (Fig 1). Under the GBR model and population growth, a KS test (marginal *α* = 0.05) cannot reject identical distributions of MAF’s of significant hits for all values of *λ* (S19 Fig). The cMR (*h* = 0) model shows an excess of intermediate frequency variants, although this does not result in rejection under the KS test (S19 Fig). If one considers trying to determine an approximate dominance coefficient in the GBR model, it would be found that there is a distribution of coefficients across sites. Yet, when simulating iMR model, we find that an intermediate degree of dominance, *h* = 0.25, results in distribution of significant hits which is similar to the GBR results (Fig 4).

We note that there is no compelling reason to expect any specific value of *λ* to be a particularly good fit to the empirical data. The empirical data are composed of genome-wide data for multiple traits. We feel that the mutational parameters, λ and mutation rate to causal variants, are likely to vary across the genome and across traits. Thus, the empirical data reflect a mixture of different underlying models and ascertainment schemes. The reason we emphasize this feature of the data is to demonstrate that models with rare alleles of large effect do not necessarily imply an observed excess of rare significant GWAS hits.

In consideration of the rare allele of large effect hypothesis, [62] proposed a model where multiple rare alleles dominate disease risk and create synthetic associations with common SNPs. However, later it was shown that this particular model was inconsistent with GWAS theoretically and empirically [4,26,74]. This inconsistency remained un-reconciled in the literature until now. We find that the MAF distribution of significant hits in a GWAS varies widely with choice of genetic model. In particular, we confirm the results of Wray et al. [26], that AC evolutionary models predict an excess of low frequency significant hits unless trait effect sizes are quite small. Also, the cMR model predicts an excess of intermediate and common significant hits. Utilizing a GBR model or an intermediate iMR model with *h* = 0.25−0.5, reconciles this inconsistency by simultaneously predicting the importance of rare alleles of large effect and the correct allele frequency distribution among statistically significant single markers.

## Conclusion

Several empirical observations provide support for the presence of gene-based recessivity underlying variation for some complex traits in humans. The minor allele frequency distribution of significant GWAS hits is relatively flat [4, 75], which our results show is consistent with either the presence of small additive effect loci or gene-/site-based partially-recessive loci with intermediate to large effects (Fig 4). Models with loci of large additive effects predict an excess of rare significant hits. Oppositely, models with complete site-based recessivity predict an excess of common significant hits for all simulated mutation effect size distributions.

SNP based estimates of dominance heritability are much lower than estimates of dominance from twins [27, 68]. Of the models we explored, only the gene-based recessive model with intermediate to large effects is consistent with difference between twin and SNP based estimates of dominance variance (Fig 2). Under a site-based recessive model of partial recessivity (*e.g.* h =0.25), there should be no significant difference between estimates of dominance variance from SNP and twin studies, provided that the statistical assumptions are met for both approaches (Fig 2). Our findings support a more thorough investigation into the importance of compound heterozygosity in the genetics complex-traits. However, it may be difficult to directly observe non-additive gene-level effects through analysis of individual SNP markers.

Additionally, the genetic model appears to be important in the design and analysis of association studies. While changes in population size do affect the relationship between effect size and mutation frequency [48–61] (Fig 1 and S5 Fig), different mappings of genotype to trait value do this in radically different ways for the same demographic history (Fig 1). From an empirical perspective, our findings suggest that re-sequencing in large samples is likely the best way forward in the face of the allelic heterogeneity imposed by the presence of rare alleles of large effect. Re-sequencing of candidate genes [76–79] and exomes [40, 80–85] in case-control panels have observed an abundance of rare variants associated with case status. Here we show that under a model of mutation-selection balance at genes, neither current single-marker nor popular multi-marker tests are especially powerful at detecting large genomic regions harboring multiple risk variants (Fig 3). However, we show that using permutations to derive p-values improves the power of SKAT [69] with a linear kernel (equivalent test statistic to c-Alpha [38]). Similarly, another permutation based test, the ESM test [72], has more robust power across demographic and genetic models (Fig 3).

Conceptually, *cis*-effects arise naturally from the original definition of a gene in which mutant recessive alleles fail to complement [35]. We show that cis-effects within a locus, represented by the GBR model, can have an important impact on the population level architecture of a complex trait. This conclusion is important for future simulation studies as well as the interpretation of empirical data. It is important to note that despite our use of the term ”gene-based” this model may apply to any functional genomic element in which there are multiple mutable sites affecting a trait in in *cis*, not just to genes. From a theoretical perspective, our work motivates the development of a more generalized gene-based model to include arbitrary dominance and arbitrary locus size. Empirically, we find that the GBR model is broadly consistent with a variety of observations from the human statistical genetics literature. Thus, there is an evident need for improved region-based association tests and the development of genetic variance component methods for haplotypes.

## Materials and Methods

### Forward simulation

Using the fwdpp template library v0.2.8 [86], we implemented a forward in time individual-based simulation of a Wright-Fisher population with mutation under the infinitely many sites model [87], recombination, and selection occurring each generation. We simulated populations of size *N =* 2e4 individuals for a time of 8*N* generations with a neutral mutation rate of *µ* = 0.00125 per gamete per generation and a per diploid per generation recombination rate of *r =* 0.00125. Deleterious mutations occurred at a rate of *µ*_*d*_ = 0.1*µ* per gamete per generation. These parameters correspond to *θ* = 4*Nµ* = *ρ* = 4*Nr* = 100 and thus our simulation approximates a 100Kb region of the human genome. For simulations with growth, we simulated an additional 500 generations of exponential growth from *N*_*i*_ = 2*e*4 to *N*_*final*_ = 1*e*6. This demographic model is much simpler than current models fit to empirical data [58]. However, this simple model allows us to more easily get a sense of the impact of population expansion [21, 22]. 250 simulation trials were performed for each parameter/model combination.

### Exploring the gene region’s contribution to heritability

Broad-sense heritability can be calculated directly from our simulated data as 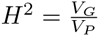. We explored broad-sense heritability as a function of mean causative effect size under each model. We compare our simulation results to 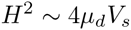 for additive models and 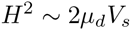 for recessive models [88, 89]. In our simulations, *V*_*s*_ = 1, and we tuned the environmental variance *V*_*e*_ to generate simulations for which *E*[*H*^2^] ~ 0.04 or ~ 0.08.

### Determining the genetic load of the population

Genetic load is defined as the relative deviation in a populations fitness from the fitness optimum, 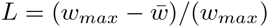. We set the phenotypic optimum to be zero; *P*_*opt*_ = 0. When determining fitness for the SBR models, we subtract one from all phenotypes. This implies that 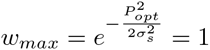 and that load is a simple function of the phenotypes of the population, 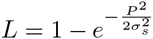. We also used the mean number of mutations per individual, and the mean frequency and effect sizes of segregating risk variants as proxies for the genetic load [21, 90].

### Additive and dominance genetic variance over allele frequency

We used an approach based sequential (type-1) regression sums of squares to estimate the contribution of the additive and dominance effects of variants to the total genetic variation due to a locus. Given a genotype matrix (rows are individuals and columns are risk variants) of (0,1, or 2) copies of a risk allele (*e.g.* all mutations affecting phenotype), we sort the columns by decreasing risk mutation frequency. Then, within frequency classes, columns were sorted by decreasing effect sizes. For each variant a dominance component was also coded as 0, 2q, or 4q-2 according to the orthogonal model of [27], where q is the frequency of the variant in the population. We then used the R package biglm [91] to regress the individual genetic values (*G* in the previous section) onto this matrix. The variance explained by the additive and dominance effects of the *m* markers with *q ≤ x* is then approximately 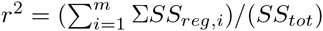. Averaging results across replicates, this procedure results in a Monte-Carlo estimate of the fraction of *V*_*G*_ that is due to additive and dominance effects of variants with population frequency less than or equal to *x* is 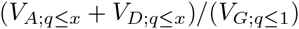 [21]. This fraction can be easily partitioned into strictly additive and dominance components.

### Additive and dominance heritability in random population samples

We employed three different SNP-based approaches to estimating heritability from population samples: GREMLd, minor allele frequency stratified(MS) GREMLd [27], and MS-Haseman-Elston (HE) regression [92, 93]. For comparison, we calculated the true total heritability in the sample as 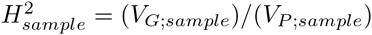. Unfortunately, due to the nature of our simulated data MS-GREMLd did not result in sufficiently reliable results. Under MS-GREMLd, many replicates resulted in numerical errors in GCTA. These problems were present at a rate of less than 1/100 replicates using non-MS GREMLd, but were increased by splitting the data into multiple GRMs.

Using raw individual phenotypic values as quantitative trait values, random samples from simulated populations (n=6000) were converted to .bed format using PLINK 1.90a [94]. PLINK was also used to test for HWE (*p* < 1*e* − 6) and filter on minor allele frequency. GCTA 1.24.4 [34] was used to make genetic relatedness matrices (GRM) for both additive and dominance components with the flags –autosome and –make-grm(-d).

For non-MS runs, we tested the effect of filtering on MAF by performing the analysis on unfiltered datasets and with markers with *MAF* < 0.01 removed. For MS estimates we stratified the additive and dominance GRM’s into two bins *MAF* ≤ 0.01 and *MAF* > 0.01. GREMLd analysis was performed in GCTA with Fisher scoring, no variance component constraint and a max of 200 iterations. MS-HE regression was carried out by regressing the off diagonal elements of each GRM onto the cross product of the scaled and centered phenotypes in a multiple linear regression setting in R [95].

### Twin studies

To simulate twin studies we sampled 2000 monozygotic (MZ) and 2000 dizygotic (DZ) twins pairs from the final generation of the simulations. Parents were sampled randomly without replacement. MZ twin pairs were formed by sampling a single gamete pair, one recombinant from each parent, and two environmental random deviates. DZ twin pairs were formed by sampling two gamete pairs, two recombinant gametes from each parent, and two environmental random deviates. Our simulated studies are ideal in that there are no correlated environmental effects, but potentially problematic due to low total heritability. We explored the use of structural equation modeling (SEM) using the package OpenMx [96], but chose to rely strictly on estimates of twin correlation obtained directly from the data. For monozygotic (MZ) twins, we used only a single child gamete pair with two unique environmental deviates. For dizygotic (DZ) twins we used two child gamete pairs, each with a unique environmental deviate. Broad sense heritability is the correlation between MZ twin pairs; 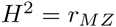. Under a purely additive model, the DZ twin correlation should be half of the MZ twin correlation. Non-additive genetic components of phenotypic variance reduce the DZ twin correlation. If all non-additive heritability is due to dominance, then the dominance heritability can be calculated as twice the difference between the MZ twin correlation and two-times the DZ twin correlation: 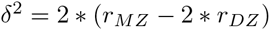. The additive heritability can then be calculated as the difference between the broad-sense and non-additive component: 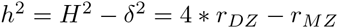 [29].

These direct estimates of MZ and DZ twin correlations in our simulations are reliable as we have no measurement error, shared environmental effects, gene-by-environment effects, or gene-by-gene interactions. Additionally, we only simulate a single genomic region contributing *H*^2^ ~ 0.04, which made use of SEM difficult numerically. This creates a limitation in that we can not discuss when a model with dominance is a better fit to the data than the additive only model. But, the benefit of using direct estimates is that we can clearly see what signals are present in the data. To further clarify the data visualization, we pooled our 512 twin-study replicates into groups of 8, creating 64 sets of MZ-DZ twin phenotypes. This did not have an effect on the central tendencies of our estimates, but it reduced the variance. The twin study error bars in Fig 2 are based on 64 sets of 64,000 individuals, which is larger than a typical twin study. However, one reason our results have high variance is because we only simulate a single locus, rather than a whole trait.

### Case-control studies

Following [36], we sampled 3000 cases and 3000 controls from each simulated population. Cases were randomly sampled from the upper 15% of phenotypic values in the population, and controls were randomly sampled from within 1 standard deviation of the population mean. This is the liability scale model (see [29]). We define a ”GWAS” to be a study including all markers with MAF ≥ 5% and a re-sequencing study to include all markers. In all cases we used a minor allele count logistic regression as the single marker test. For single marker tests, the p-value cut off for significance is *p* ≤ 1*e* — 08 which is common in current GWAS [62, 97]. Power is determined by the percentage of simulation replicates in which at least one marker reaches genome wide significance.

### Region-based tests of association due to rare alleles

We applied multiple region-based tests to our simulated data, *ESM*_*K*_ [36], several variations of SKAT [39] and c-Alpha [38]. We used the R package from the SKAT authors to implement their test (http://cran.r-project.org/web/packages/SKAT/index.html). The remaining tests were implemented in a custom R package (see Software availability below). For the *ESM*_*K*_ and c-Alpha we performed up to 2*e*6 permutations of case-control labels to determine empirical p-values. Common variants (*q* ≥ 0.05) were removed prior to performing region-based rare variant association tests.

### Distribution of Significant GWAS Hits

Following [4, 26], we calculated the distribution of the minor allele frequency (MAF) of the most significant SNPs in a GWAS in empirical and simulated data. The empirical data was obtained from the NHGRI-EBI GWAS database (http://www.ebi.ac.uk/gwas/) on 02/05/2015. We considered the same diseases and applied the same filters as in Table 3 of [26]. Specific information regarding the empirical data can be obtained in S1 Table.

In order to mimic ascertained SNP data, we sampled markers from our case/control panels according to their minor allele frequencies [98], as done in [36]. Additionally, we removed all markers with MAF < 0.01 to reflect common quality controls used in GWAS. The simulated data were grouped by genetic model, demographic scenario, heritability level, and mutation effect distribution. We then plotted the minor allele frequency of the most significant marker with a single-marker score −*log*_10_(*p*) ≥ 8, for all replicates where significant markers were present.Finally, we performed a two-sample KS test in R between each group of simulated GWAS hit allele frequencies and the empirical data.

### Human demography

We simulated a demographic model for Europeans based on [40] as described in [21]. For simplicity, we ignored migration between the European (EA) and African American (AA) populations. The model was implemented using the Python package `fwdpy` version 0.0.4, which uses `fwdpp` [86] version 0.5.1 as a C++ back-end. During the evolution of the EA population, we recorded the genetic variance in the population, *V*_*G*_, and the number of deleterious mutations per diploid (a measure of genetic load [21]) every 50 generations. In a separate set of simulations, we applied the regression method described above to calculate cumulative additive genetic variance as a function of allele frequency. Because the regressions are computationally demanding, we applied the method in the generation immediately before, and at the start of, any changes in population size.

These simulations were run with no neutral mutations, and the recombination rate and mutation rate to causative mutations were the same as in the simulations described above.

The Python scripts for these simulations and iPython/Jupyter notebooks used for generating figures are available online (see Software availability section below).

### Software availability

Our simulation code and code for downstream analyses are freely available at

- http://github.com/ThorntonLab/disease_sims
- http://github.com/molpopgen/buRden
- http://github.com/molpopgen/fwdpy
- http://github.com/molpopgen/TennessenEAonly

